# Dissecting multitrophic interactions: the relationships among *Entomophthora*, their dipteran hosts, and associated bacteria

**DOI:** 10.1101/2025.03.20.644288

**Authors:** Zuzanna Płoszka, Karol H. Nowak, Marta Tischer, Anna Michalik, Michał R. Kolasa, Piotr Łukasik

## Abstract

Interactions with microorganisms from across a range of mutualism-to-pathogenicity spectrums shape the biology of insects at all levels - from individual traits to populations and communities. However, the understanding of pathogens attacking non-model insect species in natural ecosystems, or their interactions with other insect-associated microorganisms, is fragmentary.

Here, we tested a conceptually novel approach - the simultaneous sequencing of insect, fungal, and bacterial marker gene amplicons - as a means of dissecting interactions among entomopathogenic fungi in the genus *Entomophthora* and their dipteran hosts in South Greenland. We aimed to describe the taxonomic diversity of *Entomophthora*-killed flies, their pathogens, and the bacterial diversity within a set of field-collected dead insects exhibiting signs of *Entomophthora* infection.

Across nine collected dipteran species, we identified multiple *Entomophthora* genotypes, with strong but not perfect patterns of host-specificity across the five targeted marker regions. Additionally, we found consistent differences in bacterial community composition among fungus-killed fly species and sampling sites. Our results substantially expand the knowledge of *Entomopthora* diversity and host associations while providing the very first insights into associated bacteria and their potential roles. We also conclude that multi-target amplicon sequencing can be a powerful tool for addressing broad questions about biological interactions in diverse natural communities.

## Introduction

Virtually all living organisms are embedded in complex networks of interactions with other organisms. These interactions, encompassing food organisms, mutualists, commensals, competitors, and natural enemies, determine the species’ positions and roles within the ecosystems and their evolutionary trajectories [1,2]. In the epoch of rapid environmental change and biodiversity crisis [3], reconstructing these relationships is more critical and urgent than ever. Despite that, as we struggle to catalog Earth’s biodiversity - with a mere one-fifth of predicted insect species and an even smaller fraction of fungal species ever formally named [4] - we know much less about their biological relationships. the diversity and specificity of pathogens attacking wild non-model organisms, or conversely, the range of host species suitable for a specific parasite or pathogen strain, are among those poorly understood. Such studies are particularly challenging in systems where the interacting organisms cannot be confidently assigned to species based on morphology alone, as in the case of fungal pathogens of insects [5]. The research focus on the few fungal species that can be used as biocontrol agents for agricultural pests [6,7] has left the vast majority of other insect-associated fungi largely unknown. However, new methods, especially next-generation sequencing technologies combined with improving bioinformatic tools and expanding reference databases, facilitate and speed up the efforts to characterize fungal entomopathogen diversity [8,9]. They could also help uncover other, even less understood levels of insect-fungus interactions, including how they may be influenced by their co-occurring bacteria [10,11].

Some of the most notorious insect pathogens belong to the order Entomophthorales within the phylum Zoopagomycota [12]. One of its best-known representatives is the “zombie fly” fungus, *Entomophthora muscae*, inducing “summit disease” in its dipteran hosts, where shortly before death, infected flies are induced to climb to elevated points where they attach before death, facilitating spore dispersal [13]. Interestingly, sporulating cadavers can induce strong sexual attraction in males, which facilitates fungal infection [14,15]. Using such mechanisms, these fungi can reach high infection prevalence and decimate host populations [16].

Such precise host manipulation is generally linked to pathogen specificity [17] and, thus, the anticipated high pathogen genetic diversity across taxonomically diverse hosts. Indeed, the genus *Entomophthora* is diverse, with 21 named species and substantial genetic diversity within species. Marker gene sequences amplified from different host species or genera tend to form distinct clades on the phylogenies, which corresponds to host-specificity [18]. On the other hand, laboratory experiments have revealed the ability of the same *E. muscae* strains to infect diverse Diptera [17], suggesting that host shifts may happen occasionally. However, the actual *Entomophthora* diversity and host range in diverse natural insect communities remain poorly understood [13]. Here, we hypothesized that in remote Arctic environments with low species diversity [19], likely with fewer opportunities for independent colonization by different fungal strains, fungal host ranges may be broader. Recent advantages in sequencing and bioinformatic tools have made testing such hypotheses possible.

During our recent fieldwork in South Greenland, we frequently encountered diverse dead flies with symptoms characteristic of *Entomophthora* infection (Figure 1). This included cases where on a single plant we found hundreds of individuals representing different dipteran species. We asked about the diversity and host specificity of *Entomophthora* attacking these South Greenland flies. Specifically, we aimed to: (1) validate multi-target amplicon sequencing as a tool for cost-effective characterization of host-pathogen interactions in such natural communities; (2) describe the taxonomic diversity of the dipteran hosts of *Entomophthora* in the Narsarsuaq region of South Greenland, and (3) of *Entomophthora* genetic diversity and specificity across host species. Finally, (4) we aimed to assess bacterial diversity within fungus-infected Diptera across species and sites. We addressed these goals by applying an innovative, custom approach - the simultaneous sequencing of insect, fungal, and bacterial marker region amplicons - to a collection of 77 fungus-killed Diptera.

**Figure 1:**
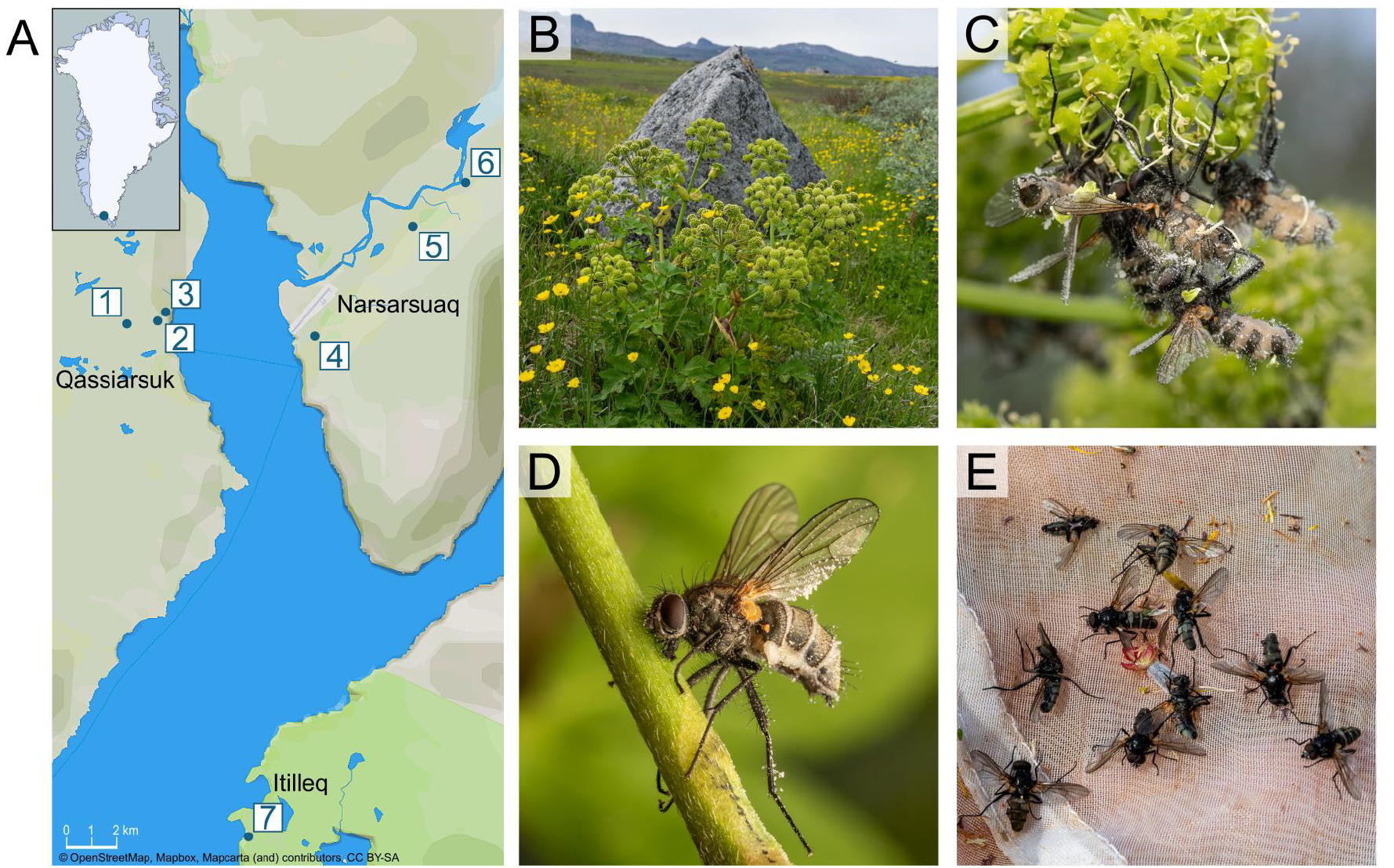
**A**. Insect sampling sites in South Greenland; **B**. Itilleq site (#7) with *Angelica archaengelica*, from where a large share of specimens originated; **C**. A cluster of *Entomophthora*-killed flies on *A. archaengelica*; **D**. *Entomophthora*-killed muscid *Thricops longipes*, the most abundant species in the collection; **E**. Dead flies displaying symptoms of *Entomophthora* infection in a sweep net sample. Photographs: Piotr Łukasik.

## Materials and methods

### Insect sampling and selection

The study material was collected within a 20-kilometer radius from the town of Narsarsuaq, Kujalleq municipality, Southern Greenland, between the 3rd and 13th of July 2023 (Figure 1A-B; Supplementary Table S2) (permit no. G23-047). While broadly sampling diverse local insects, we identified *Entomophthora*-killed flies by their characteristic posture, with splayed wings, and usually with fungal conidiophores growing out from the abdominal cuticular segments and by abundant conidia on insect bodies and adjacent surfaces [13]. We collected these insects in two ways. First, the dead flies were hand-picked from vegetation (Fig. 1C-D). This included abundant assemblages of *Entomophthora*-killed Diptera, often comprising dozens of dead insects, on *Angelica archaengelica* flower clusters (Fig. 1C). Additionally, we identified dead flies in sweep net samples: after we swept vegetation and removed living insects with an aspirator, occasionally one or more dead dipterans in a characteristic pose remained within a net and was collected (Fig. 1E). All specimens were preserved in 95% ethanol, transported to the laboratory at the Institute of Environmental Sciences of Jagiellonian University in Poland, and stored at −20°C until further processing. Finally, fungus-infected insects were pre-sorted based on morphology and representative individuals from each morphospecies were photographed under the binocular microscope. The species names were obtained using a combination of dichotomous keys [19] and COI barcodes, as described further.

### Laboratory processing

Samples were processed following a modified protocol detailed recently [20] and used in other recent projects conducted by the Symbiosis Evolution Research Group [21–23]. Briefly, individual insects were placed in 2ml screw-cap tubes with lysis buffer and ceramic beads, homogenized on Omni Bead Raptor Elite homogenizer, and then digested for two hours at 55°C. 40 µl of the homogenate was then purified using 2x volume (80 µl) of SPRI beads and then eluted in 20 µl of elution buffer.

We then prepared multi-target amplicon libraries for different combinations of insect, fungal, and bacterial marker genes. We followed a two-step-PCR library preparation protocol, where in the first step, we used a combination of template-specific primers with Illumina adapter stubs, and in the second indexing PCR step, we added a unique combination of indexes for each library and completed adapters. During each laboratory step, we added negative control samples to capture potential contamination during that step. We amplified the total of 9 marker regions from experimental samples across three experimental batches (Supplementary Table S4-12). In the first batch, for the first PCR, we combined insect mitochondrial COI primers BF3-BR2, universal primers for the fungal 18S rRNA V7-V8 and ITS2 regions, and bacterial 16S rRNA hypervariable V4 region. As the data provided virtually no phylogenetic resolution for *Entomophthora*, in the second experiment, we used newly designed *Entomophthora*-specific primers to amplify and sequence two ITS regions (ITS1 and ITS2) and three protein-coding genes: RNA polymerase II 2nd largest subunit (RPB2), minichromosome maintenance complex component 7 (MCM7) and elongation factor 1-alpha (EF1α). Because of highly uneven read numbers among regions, in the third experiment, we used the same primers as in the second (except ITS2) but amplified ITS1 in a separate reaction from three protein-coding genes before pooling the products for the indexing step. The sequences of all primers and details of reaction mixes and cycling programs are provided in Supplementary Table S1.

Final amplicon libraries were quality-verified on 2% agarose gel, pooled based on the band brightness, and sequenced on Illumina NextSeq 2000 at the Genomics Core Facility, Małopolska Centre of Biotechnology of Jagiellonian University, using 600-cycle P1 and P2 flow cells.

### Bioinformatic analysis

We followed modified bioinformatic protocols described in detail recently [21,23]. Initially, reads for each library were split into bins corresponding to different targets based on primer sequences. If the same target was amplified in different experiments, we only used the data from the last experiment. Then, for each bin, we used a custom pipeline combining custom Python scripts with already established bioinformatics tools, available on the GitHub page https://github.com/ZZPloszka/ento_amplicons. First, forward and reverse reads were quality-filtered and assembled into contigs using PEAR [24]. Next, contigs were de-replicated [25] and denoised [26] separately for every library to avoid losing information about rare genotypes during the denoising of the whole sequence set at once [27]. The sequences were then screened for chimeras using USEARCH and classified by taxonomy using the SINTAX algorithm and customized databases: SILVA (version 138 SSU) for bacterial 16S rRNA and fungal 18S rRNA [28]. The identity of *Entomophthora* protein-coding sequences was verified through BLASTn comparisons against the NCBI database. Finally, the sequences were clustered at a 97% identity level using the UPARSE-OTU algorithm implemented in USEARCH. We produced data summary tables with two levels of classification: zOTUs (zero-radius Operational Taxonomic Units) to reconstruct genotypic diversity across the collection and 97% OTUs (Operational Taxonomic Units) to assess broader patterns of bacterial clade distribution across hosts.

We also filtered the 16S rRNA dataset for putative reagent contamination using negative controls for different laboratory steps, as described recently [29,30]. Briefly, for all zOTUs, we calculated their relative abundances in all samples and then compared the maximum relative abundance values for each zOTU in experimental samples as well as in blanks. zOTUs with an experimental-to-blank ratio of less than 10 were classified as laboratory contaminants and excluded. Further, 5 samples where >70% of reads were classified as contaminants were also removed from the analysis of bacterial diversity [31]. We manually verified input and output files to ensure that no abundant genotypes were erroneously excluded during the filtering procedure.

### Statistical analysis and data visualization

For phylogenetic analyses, we used sequences of genotypes identified as Entomophthorales. Genotype sequences were obtained for five genes: ITS1, ITS2, MCM7, Ef1α, and RBP2. For each gene, we performed a separate alignment using the MAFFT v7 program [32]. Phylogenetic trees were reconstructed with a maximum likelihood approach for each gene alignment separately, using the software IQ-TREE v2.3.6 [33]. Statistical support for tree topologies was calculated by 1000 ultrafast bootstrap replicate [34] and the SH test [35] implemented in IQ-TREE. The best-fitting substitution models for tree inference were determined using the ModelFinder algorithm implemented in IQ-TREE, based on the Bayesian Information Criterion [36] (for ITS1 best-fit model was HKY+F, for ITS2 K3Pu+F, for MCM7 TN+F, for Ef1a K2P+I, and for rbp2 TNe+I). Phylogenetic trees were visualized with iTol Interactive Tree of Life (iTOL) v6 [37].

To assess differences in community composition among populations, we performed a Permutational Multivariate Analysis of Variance (PERMANOVA) [38] using the adonis2 function from the vegan package [39] in R. The analysis was based on a Bray-Curtis distance matrix [40] and tested the effect of population on community composition. We used 999 permutations to assess statistical significance.

Data were visualized using custom scripts written in Processing 3.5.4 [41], available in the GitHub repository (https://github.com/ZZPloszka/ento_amplicons).

### Data availability

Raw amplicon sequencing data have been deposited in NCBI’s Short Read Archive (BioProject Accession no. PRJNA1214689). The reconstructed insect and fungal marker-gene sequences are available in Supplementary Tables S4-12.

## Results

### Validation of multi-target amplicon sequencing for high-throughput characterization of host-pathogen interactions

Our custom amplicon sequencing-based characterization of insect-entomopathogen-bacteria interactions was developed in several steps. Initially, we targeted a set of relatively standard markers that were simultaneously amplified from the same insects: insect COI, fungal ITS2 and 18S rRNA, and bacterial 16S rRNA (Tables S4-7, Fig 2A). COI data allowed us to identify most specimens (77 out of 86) to species level, despite the inconsistent numbers of reads (Supplementary Tables S3-S4), while 16S rRNA data provided biologically realistic information about bacterial community composition across most samples (see below). Unexpectedly, *Entomophthora* was virtually absent from ITS2 data: the reconstructed genotypes represented diverse other fungi, plus other organisms such as *Angelica archangelica* plant that many specimens were collected from (Fig. 2A; Table S7). In contrast, the vast majority of 18S rRNA reads represented *Entomophthora*: three dominant sequence variants (zOTUs) made up 98.4% of all reads, with two of these variants only present in one of the species (Fig. 2A, Table S6).

**Figure 2.**
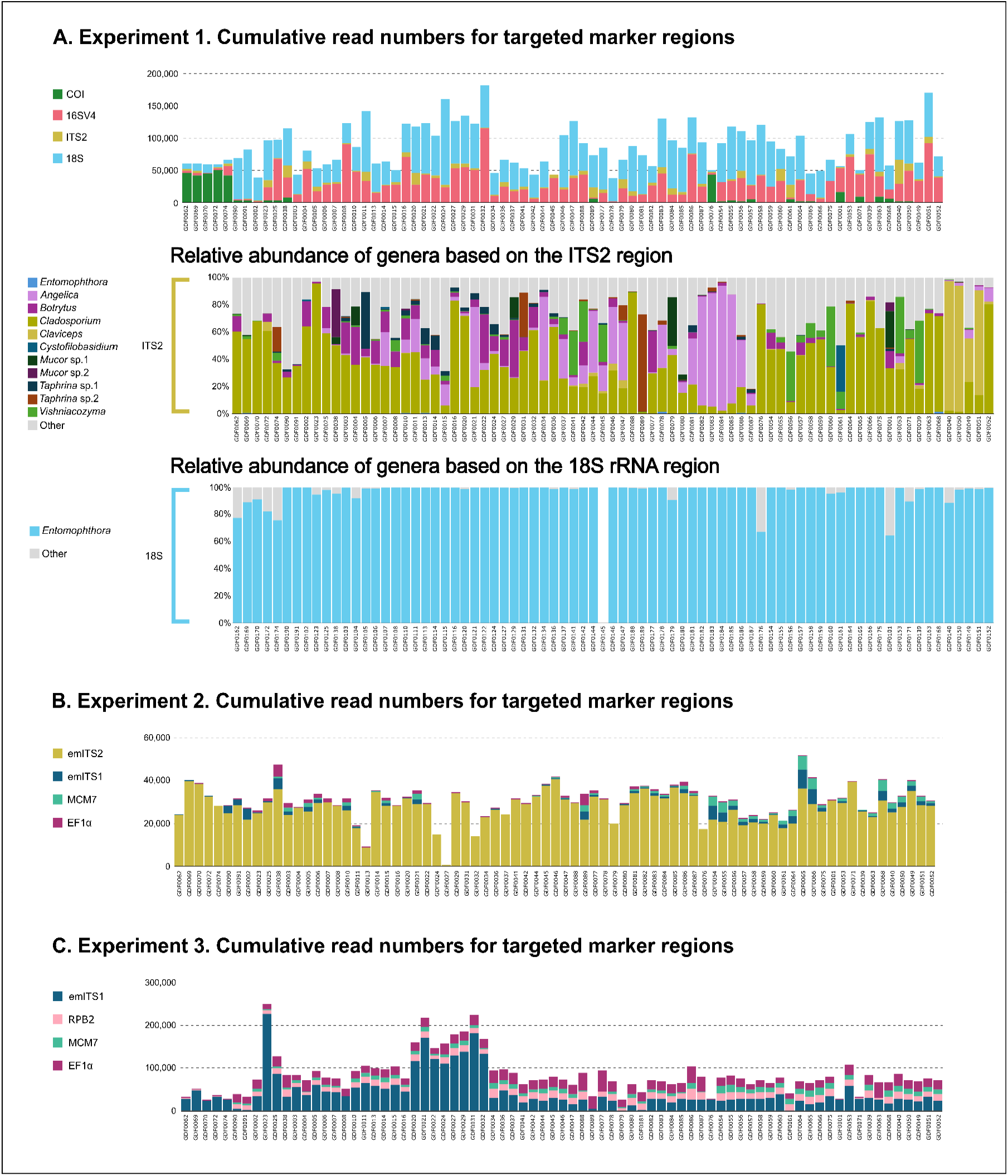
**A**. Numbers of amplicon sequencing reads corresponding to different marker regions for 77 specimens of *Entomophthora*-killed Diptera amplified in Experiment 1, and the relative abundance of fungal genera reconstructed based on two regions targeted in that experiment. **B-C**. Numbers of sequencing reads corresponding to amplified marker regions in Experiments 2-3, respectively.

The realization that the entire *Entomophthora* ITS2 region may be too long for amplification with standard primers and characterization using the Illumina platform [42] prompted us to develop new *Entomophthora*-specific primers for a portion of that region (Table S1). At the same time, to increase phylogenetic resolution, we developed new *Entomophthora*-specific primers for the ITS1 region and for protein-coding genes often used for fungal phylogenetics research: MCM7 [43], RPB2, and EF1α (Table S1) [44]. In the second experiment, we successfully amplified and sequenced these five targeted fungal regions, but the large majority of all reads corresponded to only one of these regions (ITS2), with the proportions of reads for other regions uneven and several sample-region combinations lacking (Fig. 2B). Hence, in the third batch, we adjusted primer concentrations and separately amplified (i) ITS1, and (ii) three protein-coding genes, before combining the products for the indexing step. This approach provided data for all targeted fungal regions for all samples (Fig 2E). Ultimately, across three rounds of library preparation, we have successfully amplified all targeted regions for the large majority of libraries, verifying that the results agreed with biological expectations.

### Taxonomic diversity of *Entomophthora*-killed flies

In Experiment 1, we obtained between 2 and 49783 COI reads (median: 195, average: 4537,6) for 77 individual flies (Table S4), excluding specimens where COI data did not yield products and where we lacked confidence about morphology-based identifications. Most of these reads (98.4% in a library on average) represented Diptera, and in all specimens, one COI zOTU, designated as the barcode, was represented by many more reads than others (Table S4). The 15 distinct barcode sequences identified from across 77 processed insects were confidently assigned to nine different dipteran species representing six families: *Thricops longipes* and *Lispe pumila* (Muscidae), *Botanophila rubrigena* (Anthomyiidae), *Scathophaga furcata* and *S. litorea* (Scathophagidae), *Dolichopus plumipes* and *D. groenlandicus* (Dolichopodidae), *Platycheirus hyperboreus* (Syrphidae) and *Allopiophila vulgaris* (Piophilidae) (Fig. 3). These COI-based identifications overlapped with morphology-based genus-level identifications, and allowed us to distinguish among two distinct species in the genus *Dolichopus*.

**Figure 3.**
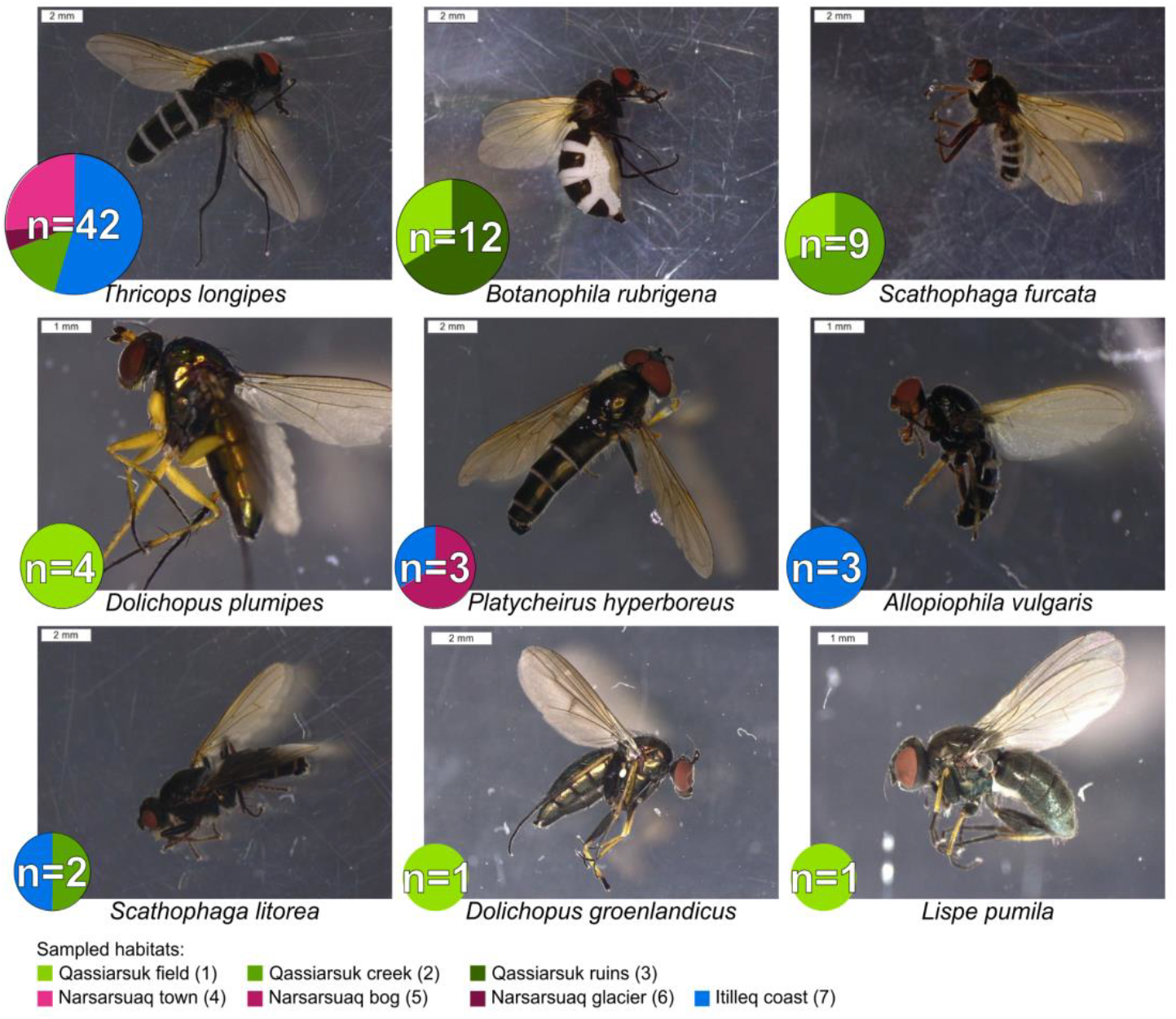
Representative specimens of nine *Entomophthora*-killed dipteran species that we encountered in South Greenland. Information about the number of characterized specimens is written into pie charts, which also indicate specific sampling sites (see Fig. 1).

Dipteran species varied in the number of specimens characterized. The most abundant, *Thricops longipes*, was represented by 42 individuals, but six species were represented by not more than four individuals. Species also varied in their distributions across collection sites and methods (Fig.3, Tables S2-S3). For example, virtually all processed *T. longiceps* specimens were hand-collected from *Angelica* plants in three distinct locations, while several other species were collected through sweep-netting of diverse plant assemblages at other locations.

### Reconstruction of *Entomophthora* diversity and distribution based on different marker regions

For the five *Entomophthora* marker regions, ITS1, ITS2, MCM7, RPB2 and EF1α we obtained an average of 43,793, 27,479, 7,350, 12,408 and 18,889 reads per specimen, respectively. For each of the regions, we observed several genotypes, abundant in multiple specimens, that had >90% sequence identity to published *Entomophthora* references (Fig. 4).

**Figure 4:**
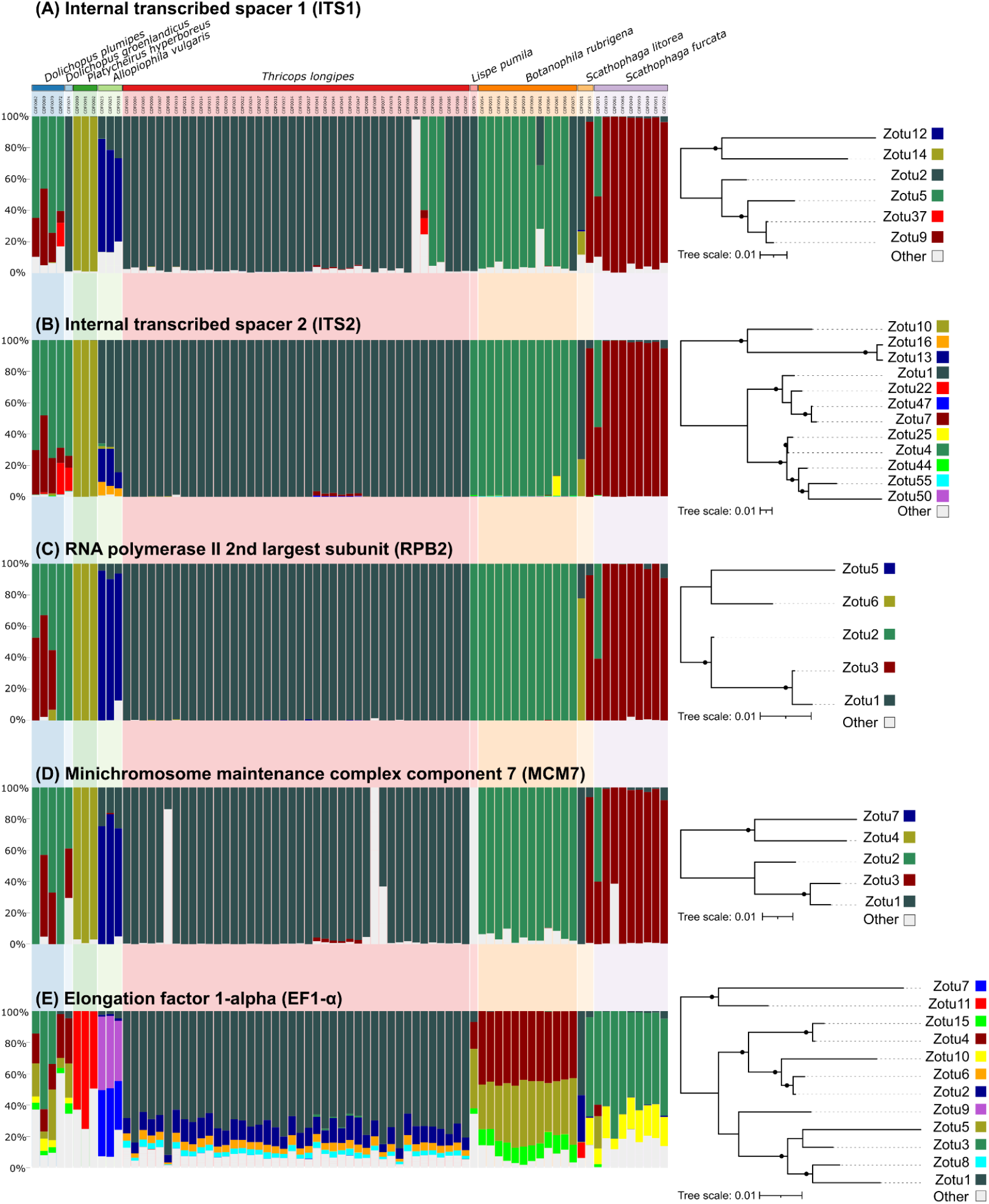
The relative abundance of amplified sequence variants of five *Entomophthora* marker regions in 77 individual flies representing nine species. Colors indicating distinct variants in different panels were selected to highlight the similarity in variant distribution patterns among marker regions. In phylogenies for each region, black dots indicate bootstrap support values above 50%.

For four of the regions (ITS1, ITS2, RPB2, MCM7 - Fig. 4A-D; Supplementary Tables S8-S11), individual insects typically carried one abundant sequence variant (zOTU). However, a substantial share of specimens had more variants - up to 4 per region. With few exceptions, the distribution of these fungal genotypes across species was highly specific. For example, the large majority of *T. longipes* individuals carried the same single variant at each of the regions, although a fraction carried a distinct or more diverse profile for one of the regions, but typically not for others. Likewise, each of the *P. hyperboreus* individuals carried the same sequence variant for each of these four regions — and they were all distinct from variants carried by any other fly. On the other hand, *A. vulgaris* and *D. plumipes* individuals carried a mixture of up to 4 variants for each of the regions, but different individuals of either species had the same set of variants. Some of the sequence variants overlapped among host species. There were some exceptions to the specificity, though. In particular, a single *S. furcata* specimen, GDF071, carried the same combination of variants across all marker regions as most individuals of *D. plumipes* - standing out from the remaining *S. furcata* individuals. Two *S. litorea* individuals carried combinations of variants very distinct from each other; one matched the dominant pathogen of *S. furcata*, while the other was unique. Likewise, *Dolichopus* individuals differed somewhat in the exact sets of variants carried.

The taxonomic classification of ITS2 sequence variants confidently assigned all abundant variants to the genus *Entomophthora*. Variants from a syrphid *P. hyperboreus* and two *Scathophaga* species were confidently classified as *E. syrphi* and *E. scathophagae*, respectively; others were classified with low confidence as *E. muscae* or matched to unclassified strains (Supplementary Table S9). Sequences for other regions generally had >95% nucleotide identity to some of the *Entomophthora* sequences in NCBI databases (Fig. 4).

Phylogenetic analysis of reconstructed sequences for these four regions highlighted the genetic variation in our dataset (Fig. 4A-D). Because of the limited length of the targeted regions, the resolution of the reconstructed phylogenetic trees was limited, and the support values for the nodes were generally low. Despite this, we can see similar patterns among trees for different marker regions. For example, sequence variants from *P. hyperboreus* (golden) and the dominant genotype from *A. vulgaris* (navy blue) were consistently placed away from other genotypes (albeit not very close to each other).

The patterns for the fifth targeted region, EF1α, were different, in that each individual carried a mixture of variants (Fig. 4E, Supplementary Table S12). However, the set of variants was largely the same for each host species, with little overlap among species. The phylogenetic analysis of these sequences showed that there are two relatively divergent variants in each insect, often accompanied by an additional variant more similar to the two.

### Comparison of bacterial diversity within fungus-infected Diptera across species and sites

After filtering reagent contamination based on negative control samples, we retained an average of 29,480 (range: 663-113,974) reads per sample (Table S5), or 86% of the pre-filtering number. We excluded five specimens (four *P. hyperboreus* and one *L. pumila*) from the analysis because >70% of their bacterial reads were classified as reagent-derived contaminants, indicating a low overall abundance of bacteria. Across the 72 retained fungus-killed flies representing the remaining seven species, we identified diverse bacteria, primarily representing Gammaproteobacteria, Alphaproteobacteria, and Bacilli. Together, the 16 most abundant 97% OTUs, each representing at least 0.5% of reads in a library on average, comprised typically 90.8% of total reads per library (Fig. 5).

**Figure 5.**
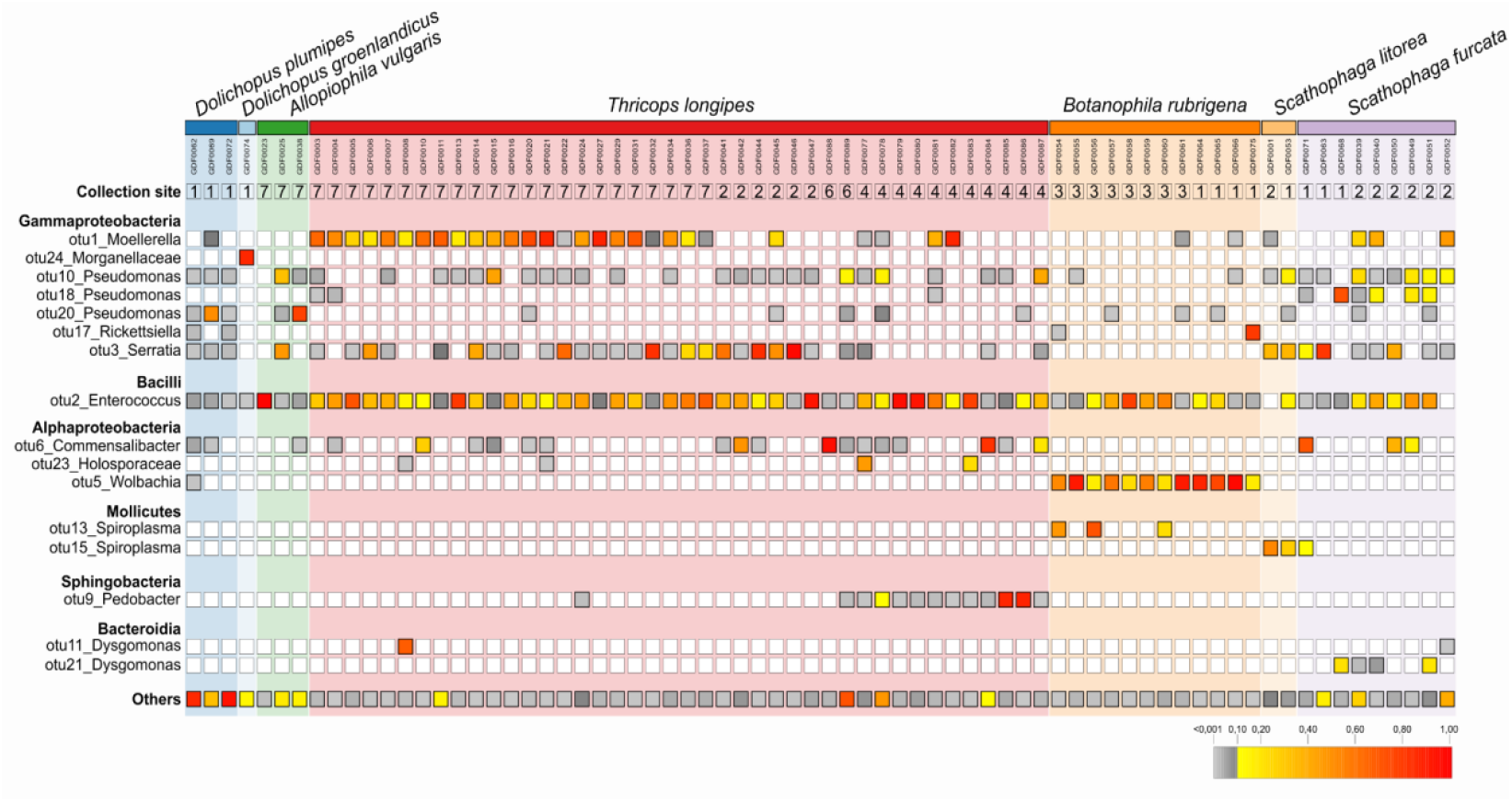
The presence and relative abundance of the 16 most abundant bacterial OTUs in *Entomophthora*-killed insects. Data for a given OTU is shown only where the relative abundance in an individual is at least 0.001, helping reduce the visual impact of potential cross-contamination among samples.

The most abundant bacterial OTU was *Enterococcus*, comprising 27% of reads in a library on average and detected in 70 specimens (97% of those processed) representing all species. Other abundant OTUs were more host species-specific. *Moellerella, Serratia*, and *Pedobacter* were present in many *T. longipes* specimens but were less common or absent in other hosts. *Wolbachia* was abundant in all specimens of *Botanophila rubrigena* and virtually absent from other species. In turn, three *Pseudomonas* OTUs were present in a large share of individuals representing most species, but their relative abundance varied among specimens and species. PERMANOVA revealed significant differences in microbial community composition among species of *Entomophthora*-killed flies (F_6,65_=3.94, p<0.001), with species explaining 26.7% of the variance in the model.

We also observed differences in bacterial associations among sites. This was most evident in *T. longipes*, for which the greatest number of samples was available. For example, *T. longipes* cadavers from Itilleq (site 7) consistently carried *Moellerella*, which was much less common at other sites, and those from Narsarsuaq town (site 4) carried *Pedobacter*, which was virtually absent elsewhere. PERMANOVA revealed significant differences in bacterial community composition among collection sites for this species (F_2,39_=4.2352, p=0.001).

## Discussion

Applying an innovative molecular approach to a collection of pathogen-killed insects from a remote area has resulted in some of the most comprehensive insights into host-entomopathogen-bacteria associations of wild insects to date. We were able to taxonomically classify the large majority of the collected insects, reconstruct biologically realistic patterns of their pathogens’ specificity, and obtain preliminary insights into bacterial communities associated with these *Entomophthora*-killed flies. These results expand our understanding of interspecific interactions, with relevance well beyond Greenland. Further, they show that the multi-target amplicon sequencing approach is an effective tool for the study of poorly characterized multi-partner interactions, with promising applications in future research. We discuss these findings and conclusions below.

### The diversity of dipteran hosts and their *Entomophthora* pathogens

*Entomophthora* and related fungi attack many diverse insects [45], but only a limited range of strains from a few host species have been systematically characterized using sequencing-based and experimental approaches [18], with a strong bias in sampling towards Europe and Eastern U.S. [13]. Here, in a geographic area with very few records to date, we have validated infections in several insect clades either not reported as *Entomophthora* hosts before or lacking host and pathogen taxonomic details. Our data provide some of the very first molecular records for *Entomophthora* colonizing several insect families and genera - as well as for their hosts. But while our collection spans some 2% of several hundred Brachycera species ever reported from Greenland [19], and likely a much larger share of species present in the Narsarsuaq area, we expect that the range of attacked clades and species is much greater than what our incidental sampling revealed.

New molecular records for *Entomophthora* genes expand our knowledge of the diversity of this important entomopathogen [13]. Sequences for some of the identified strains match established fungal species, *E. syrphi* and *E. scathophagae*, and they were discovered within the dipteran clades known to be colonized by these pathogen species [13,46,47]. Our other strains appear to represent the broader *E. muscae* species complex - a wider assemblage of genetically distinct lines that are difficult to distinguish morphologically and with scarce molecular data [46]. We assume that the majority of strains that we identified represent clades, or putative species, within the *E. muscae* species complex.

To delimit species, we require greater phylogenetic resolution than that afforded by short, conserved marker regions. We hoped to achieve this using a combination of single-copy protein-coding genes commonly used in fungal phylogenetic analyses [48,49]. However, in many specimen-gene combinations, we found more than one variant of the fungal gene sequence. This was particularly evident for EF1α (Fig. 4E), which was represented by more than one copy within two published *Entomophthora muscae* genomic assemblies. In one of the published genomes (assembly GCA_014839935.2), a search using the reconstructed EF1α sequences revealed four regions per genome that had 100% sequence coverage and >90% identity to query sequences. These patterns suggest that this gene may have existed in distinct copies already in an ancestor of the studied *Entomophthora* strains. Two other targeted protein-coding genes were present in these assemblies as a single copy or as identical copies. However, the consistent presence of the same RPB2 and MCM7 variants in replicate specimens of *D. plumipes* and *P. hyperboreus* suggests that in at least some cases, there is sequence variation within *Entomophthora* genomes. This is expected, considering that these genomes’ large size is due to repeated sequence duplications [50]. On the other hand, in some other cases, we cannot exclude the possibility of double infection of the same specimen by different fungal strains. This within-specimen sequence diversity prevented the phylogenetic analyses based on the concatenated genes that we originally planned. These data also suggest that the strategies for the phylogenomic characterization of these fungi would need to be carefully evaluated.

In previous studies, strains of *Entomophthora muscae* originating from different dipteran species or genera were found to form host-specific clades [51], despite their ability to occasionally colonize other species [18,47]. These past reports agree with the patterns encountered within South Greenland. There is no evidence supporting our hypothesis that limited dipteran diversity results in greater host ranges due to recent expansions or host shifts. However, more systematic testing of Entomophthora from across host taxonomic ranges, geographic areas, and time points is needed to address this hypothesis comprehensively.

### Bacterial diversity within fungus-infected Diptera

Interactions with bacteria are an important part of fungal biology [52] - and in the case of fungi that are pathogenic to other organisms, they are known to significantly influence the outcome of infection [53]. However, outside of a single report on *Serratia* isolated from *Entomophthora*-killed *Musca domestica* [54], we are not aware of prior efforts to systematically characterize the bacterial component of these interactions. Then, our results on bacterial communities of *Entomophthora*-killed Greenland flies - including differences among species as well as among locations - provide a completely new perspective on this system.

An intriguing question is about the origin of bacteria in the processed samples. We think that they likely come from several sources. First, the known variation among insect species in microbial community abundance and composition [55], certainly contributes to the variation that we observed. In particular, *Wolbachia* - a specialized endosymbiotic bacterium that colonizes about half of all insect species and some other invertebrates - is known to vary in prevalence within and among host populations and species [56], and its detection in all individuals of *B. rubrigena* but not in other species almost certainly reflects microbiota composition before *Entomophthora* infection. On the other hand, some bacteria may be specifically associated with *Entomophthora*, spreading alongside the fungus. Such fungus-symbiotic bacteria, often with important effects on their hosts, have been reported from several fungal clades [57,58]. The near-universal presence and high abundance of *Enterococcus* across dead flies, combined with some differences among species in zOTU distribution (Table S5), agrees with expectations for this type of association. On the other hand, some broadly distributed environmental microbes may thrive in the specific environment of *Entomophthora-*killed insect cadavers. We think that bacteria such as *Moellerella* or *Pedobacter*, both much more prevalent in some sampling sites for the abundant species *T. longipes* than in others, may represent this category. However, distinguishing among these scenarios would require a systematic comparison among insects at different stages of entomopathogen infection, including also uninfected individuals.

Additionally, it would be useful to assess the significance of these bacterial communities associated with *Entomophthora-*killed insects. On the one hand, diverse microbes within insects can protect their hosts against Entomophthorales [59]. Such entomopathogen-protective effects were demonstrated, for example, for some strains of *Wolbachia* [59,60]. While in our study, *B. rubrigena* did not seem to benefit from *Wolbachia*-conferred protection, it is plausible that the absence of *Wolbachia*, or other abundant facultative endosymbionts, in other species might prevent infection. On the other hand, symbionts of fungi may help them overcome host defenses or provide access to otherwise unavailable resources [53]. Testing these ideas and dissecting the nature of insect-fungal-bacterial associations would however require dedicated experimental studies. Furthermore, we know that other players exist in the system, including fungus-associated *Entomophthovirus* mycoviruses [61], certain to further influence these interactions.

### Broad sampling and multi-target amplicon sequencing as effective tools for studying interspecific interactions

High-throughput sequencing (HTS)-based characterization of insect marker genes - barcoding - has proven to be a particularly effective tool for characterizing wild insect diversity and distribution, revolutionizing the study and monitoring of insect communities [62]. Likewise, amplicon sequencing of standard bacterial or fungal marker regions has revolutionized the understanding of microbiota patterns and roles [63]. While not often done, combining insect and bacterial marker gene amplicon sequencing has proven to be a powerful approach for the study of biological interactions in diverse natural communities [21,23]. On the other hand, HTS-based characterization of multiple phylogenetically informative amplified marker regions has rarely been applied for the study of specific host-associated microbes: whole-genome approaches have made once-popular Sanger sequencing-based Multi-Locus Sequence Typing (MLST) schemes largely obsolete [64,65]. However, due to its costs and limited throughput, genomics is not an ideal tool for large-scale ecological surveys of host-associated or environmental microbes, especially those with large genomes.

Here, targeting phylogenetically informative *Entomophthora* genes using multiplexed custom primers, developed based on publicly available sequence resources, proved to be an effective strategy for reconstructing their diversity and specificity across dozens of field-collected dead dipterans. This cost-effective approach provided reliable data for the targeted fungal regions, even when multiple sequence variants were present in the same sample. Besides, it provided information on associated organisms - in this case, insects and bacteria. We conclude that even in the era of genomics, carefully designed multitarget amplicon sequencing can be a highly cost-effective strategy for addressing broad questions about biological interactions [23,66], easily scalable for much greater sample collections than those used here.

On the other hand, our experiences show that the implementation of such multi-target approaches is not trivial. The design and testing of primers for an organism of interest require reference sequence datasets and certain expertise, and the simultaneous amplification of different marker regions in a single multiplex reaction will likely require optimization. Here, differences in the abundance of alternative targets within samples were the likely reason for the large overrepresentation of ITS products, complicating our attempts to simultaneously characterize fungal and insect genes [67]. At the validation stage, besides adjusting primer concentrations, it may be worth considering solutions such as the separate amplification of alternative products, as we did in Experiment 3. Despite the effort required for optimization, implementing such assays may be a worthwhile investment for projects addressing broad questions about interspecific interactions in nature.

### Conclusions

Across multiple Diptera-*Entomophthora* associations that we report from a so far uncharacterized corner of the world, we identify entomopathogen distribution and specificity patterns that largely agree with expectations based on limited experimental data from other regions. While doing so, we also showcase how modern sequencing-based approaches enable cost-effective high-throughput characterization of multi-partner associations involving insects, their pathogens, and bacteria. Such interactions can dramatically affect insect populations, with cascading effects on other species and, consequently, multi-species communities [66]. With the rapid environmental changes driving global biodiversity declines and species’ range changes [3], knowing bacterial and fungal roles in shaping interaction networks is critical for understanding, predicting, and counteracting the negative processes. Our study highlights a strategy that, when implemented broadly, can revolutionize our understanding of how pathogens and protective symbionts shape natural populations and communities.

## Acknowledgments

*This project was supported by the Polish National Agency for Academic Exchange grant no. PPN/PPO/2018/1/00015 and Polish National Science Centre grant 2021/43/B/NZ8/03376, as well as a grant from the Priority Research Area BioS under the Strategic Programme Excellence Initiative at Jagiellonian University*.

## Supplementary table legends

S1_Primer_seqs - list of used primers along with their sequences, PCR setup, and PCR cycling conditions

S2_Collections - list of collection samples with descriptions, coordinates, and descriptions of sites and dates of collection

S3_Specimens - list of all specimens with unique IDs, collection sites, morphology and COI-based taxonomy, COI barcode sequences, and numbers of amplicon reads in each targeted marker region

S4_COI_zOTUs - table of COI zOTUs with taxonomic assignment and numbers of amplicon reads for each insect sample

S5_16S_zOTUs - table of 16S (bacterial) zOTUs with taxonomic assignment and numbers of amplicon reads for each insect sample

S6_18S_zOTUs - table of 18S (fungal) zOTUs with taxonomic assignment and numbers of amplicon reads for each insect sample

S7_ITS2_zOTUs - table of ITS2 (fungal) zOTUs with taxonomic assignment and numbers of amplicon reads for each insect sample

S8_emITS1_zOTUs - table of *Entomophthora*-specific ITS1 zOTUs with taxonomic assignment and numbers of amplicon reads for each insect sample

S9_emITS2_zOTUs - table of *Entomophthora*-specific ITS2 zOTUs with taxonomic assignment and numbers of amplicon reads for each insect sample

S10_emMCM7_zOTUs - table of *Entomophthora*-specific MCM7 zOTUs with taxonomic assignment and numbers of amplicon reads for each insect sample

S11_emEF1a_zOTUs - table of *Entomophthora*-specific EF1α zOTUs with taxonomic assignment and numbers of amplicon reads for each insect sample

S12_emRPB2_zOTUs - table of *Entomophthora*-specific RPB2 zOTUs with taxonomic assignment and numbers of amplicon reads for each insect sample

## References

1. McFall-Ngai M et al. 2013 Animals in a bacterial world, a new imperative for the life sciences. Proc. Natl. Acad. Sci. 110, 3229–3236. (doi:10.1073/pnas.1218525110)

2. Sudakaran S, Kost C, Kaltenpoth M. 2017 Symbiont acquisition and replacement as a source of ecological innovation. Trends Microbiol. 25, 375–390. (doi:10.1016/j.tim.2017.02.014)

3. Wagner DL, Grames EM, Forister ML, Berenbaum MR, Stopak D. 2021 Insect decline in the Anthropocene: Death by a thousand cuts. Proc. Natl. Acad. Sci. 118, e2023989118. (doi:10.1073/pnas.2023989118)

4. Hawksworth DL, Lücking R. 2017 Fungal diversity revisited: 2.2 to 3.8 million species. In The Fungal Kingdom (eds J Heitman, BJ Howlett, PW Crous, EH Stukenbrock, TY James, NAR Gow), pp. 79–95. Washington, DC, USA: ASM Press. (doi:10.1128/9781555819583.ch4)

5. Haelewaters D, De Kesel A, Pfister DH. 2018 Integrative taxonomy reveals hidden species within a common fungal parasite of ladybirds. Sci. Rep. 8, 15966. (doi:10.1038/s41598-018-34319-5)

6. Lovett B, Leger RJSt. 2017 The Insect Pathogens. In The Fungal Kingdom, pp. 923–943. John Wiley & Sons, Ltd. (doi:10.1128/9781555819583.ch45)

7. Islam W et al. 2021 Insect-fungal-interactions: A detailed review on entomopathogenic fungi pathogenicity to combat insect pests. Microb. Pathog. 159, 105122. (doi:10.1016/j.micpath.2021.105122)

8. Wu B, Hussain M, Zhang W, Stadler M, Liu X, Xiang M. 2019 Current insights into fungal species diversity and perspective on naming the environmental DNA sequences of fungi. Mycology 10, 127–140. (doi:10.1080/21501203.2019.1614106)

9. De Wint FC et al. 2024 Introducing a global database of entomopathogenic fungi and their host associations. Sci. Data 11, 1418. (doi:10.1038/s41597-024-04103-4)

10. Deng J, Xu W, Lv G, Yuan H, Zhang Q-H, Wickham JD, Xu L, Zhang L. 2022 Associated bacteria of a pine sawyer beetle confer resistance to entomopathogenic fungi via fungal growth inhibition. Environ. Microbiome 17, 47. (doi:10.1186/s40793-022-00443-z)

11. Hong S, Sun Y, Sun D, Wang C. 2022 Microbiome assembly on Drosophila body surfaces benefits the flies to combat fungal infections. iScience 25, 104408. (doi:10.1016/j.isci.2022.104408)

12. Lovett B, Macias A, Stajich JE, Cooley J, Eilenberg J, Licht HH de F, Kasson MT. 2020 Behavioral betrayal: How select fungal parasites enlist living insects to do their bidding. PLOS Pathog. 16, e1008598. (doi:10.1371/journal.ppat.1008598)

13. Elya C, De Fine Licht HH. 2021 The genus Entomophthora: bringing the insect destroyers into the twenty-first century. IMA Fungus 12, 34. (doi:10.1186/s43008-021-00084-w)

14. Zurek L, Wes Watson D, Krasnoff SB, Schal C. 2002 Effect of the entomopathogenic fungus, Entomophthora muscae (Zygomycetes: Entomophthoraceae), on sex pheromone and other cuticular hydrocarbons of the house fly, Musca domestica. J. Invertebr. Pathol. 80, 171–176. (doi:10.1016/S0022-2011(02)00109-X)

15. Naundrup A, Bohman B, Kwadha CA, Jensen AB, Becher PG, De Fine Licht HH. 2022 Pathogenic fungus uses volatiles to entice male flies into fatal matings with infected female cadavers. ISME J. 16, 2388–2397. (doi:10.1038/s41396-022-01284-x)

16. Wilding N, Perry JN. 1980 Studies on Entomophthora in populations of Aphis fabae on field beans. Ann. Appl. Biol. 94, 367–378. (doi:10.1111/j.1744-7348.1980.tb03952.x)

17. Steinkraus DC, Kramer JP. 1987 Susceptibility of sixteen species of Diptera to the fungal pathogen Entomophthora muscae (Zygomycetes: Entomophthoraceae). Mycopathologia 100, 55–63. (doi:10.1007/BF00769569)

18. Gryganskyi AP, Humber RA, Stajich JE, Mullens B, Anishchenko IM, Vilgalys R. 2013 Sequential utilization of hosts from different fly families by genetically distinct, sympatric populations within the Entomophthora muscae species complex. PloS One 8, e71168. (doi:10.1371/journal.pone.0071168)

19. Böcher J, Kristensen (†) NP, Pape T, Vilhelmsen L, editors. 2015 The Greenland Entomofauna. Brill. (ISBN:978-90-04-25640-8)

20. Buczek M, Prus-Frankowska M, Łukasik P. 2024 Quantitative multi-target amplicon sequencing workflow. (doi:10.17504/protocols.io.36wgq351ylk5/v1)

21. Kolasa M, Kajtoch Ł, Michalik A, Maryańska-Nadachowska A, Łukasik P. 2023 Till evolution do us part: The diversity of symbiotic associations across populations of Philaenus spittlebugs. Environ. Microbiol. 25, 2431–2446. (doi:10.1111/1462-2920.16473)

22. Andriienko V, Buczek M, Meier R, Srivathsan A, Łukasik P, Kolasa MR. 2024 Implementing high-throughput insect barcoding in microbiome studies: impact of non-destructive DNA extraction on microbiome reconstruction. PeerJ 12, e18025. (doi:10.7717/peerj.18025)

23. Nowak KH, Hartop E, Prus-Frankowska M, Buczek M, Kolasa M, Roslin T, Ovaskainen OT, Łukasik P. 2025 What lurks in the dark? - An innovative framework for studying diverse wild insect microbiota. bioRxiv 2024.08.26.609658. (doi:10.1101/2024.08.26.609658)

24. Zhang J, Kobert K, Flouri T, Stamatakis A. 2014 PEAR: a fast and accurate Illumina Paired-End reAd mergeR. Bioinforma. Oxf. Engl. 30, 614–620. (doi:10.1093/bioinformatics/btt593)

25. Rognes T, Flouri T, Nichols B, Quince C, Mahé F. 2016 VSEARCH: a versatile open source tool for metagenomics. PeerJ 4, e2584. (doi:10.7717/peerj.2584)

26. Edgar RC. 2016 UNOISE2: improved error-correction for Illumina 16S and ITS amplicon sequencing. (doi:10.1101/081257)

27. Prodan A, Tremaroli V, Brolin H, Zwinderman AH, Nieuwdorp M, Levin E. 2020 Comparing bioinformatic pipelines for microbial 16S rRNA amplicon sequencing. PLOS ONE 15, e0227434. (doi:10.1371/journal.pone.0227434)

28. Quast C, Pruesse E, Yilmaz P, Gerken J, Schweer T, Yarza P, Peplies J, Glöckner FO. 2013 The SILVA ribosomal RNA gene database project: improved data processing and web-based tools. Nucleic Acids Res. 41, D590–596. (doi:10.1093/nar/gks1219)

29. Mulio SÅ, Zwolińska A, Klejdysz T, Prus-Frankowska M, Michalik A, Kolasa M, Łukasik P. 2024 Limited variation in microbial communities across populations of Macrosteles leafhoppers (Hemiptera: Cicadellidae). Environ. Microbiol. Rep. 16, e13279. (doi:10.1111/1758-2229.13279)

30. Surmacz B, Stec D, Prus-Frankowska M, Buczek M, Michalczyk Ł, Łukasik P. 2024 Pinpointing the microbiota of tardigrades: What is really there? Environ. Microbiol. 26, e16659. (doi:10.1111/1462-2920.16659)

31. Łukasik P, Newton JA, Sanders JG, Hu Y, Moreau CS, Kronauer DJC, O’Donnell S, Koga R, Russell JA. 2017 The structured diversity of specialized gut symbionts of the New World army ants. Mol. Ecol. 26, 3808–3825. (doi:10.1111/mec.14140)

32. Katoh K, Standley DM. 2013 MAFFT multiple sequence alignment software version 7: improvements in performance and usability. Mol. Biol. Evol. 30, 772–780. (doi:10.1093/molbev/mst010)

33. Nguyen L-T, Schmidt HA, von Haeseler A, Minh BQ. 2015 IQ-TREE: a fast and effective stochastic algorithm for estimating maximum-likelihood phylogenies. Mol. Biol. Evol. 32, 268–274. (doi:10.1093/molbev/msu300)

34. Minh BQ, Nguyen MAT, von Haeseler A. 2013 Ultrafast approximation for phylogenetic bootstrap. Mol. Biol. Evol. 30, 1188–1195. (doi:10.1093/molbev/mst024)

35. Shimodaira H, Hasegawa M. 1999 Multiple comparisons of log-likelihoods with applications to phylogenetic inference. Mol. Biol. Evol. 16, 1114. (doi:10.1093/oxfordjournals.molbev.a026201)

36. Kalyaanamoorthy S, Minh BQ, Wong TKF, von Haeseler A, Jermiin LS. 2017 ModelFinder: fast model selection for accurate phylogenetic estimates. Nat. Methods 14, 587–589. (doi:10.1038/nmeth.4285)

37. Letunic I, Bork P. 2024 Interactive Tree of Life (iTOL) v6: recent updates to the phylogenetic tree display and annotation tool. Nucleic Acids Res. 52, W78–W82. (doi:10.1093/nar/gkae268)

38. Anderson MJ. 2017 Permutational Multivariate Analysis of Variance (PERMANOVA). In Wiley StatsRef: Statistics Reference Online, pp. 1–15. John Wiley & Sons, Ltd. (doi:10.1002/9781118445112.stat07841)

39. Oksanen J et al. 2001 vegan: Community Ecology Package., 2. 6–10. (doi:10.32614/CRAN.package.vegan)

40. Beals EW. 1984 Bray-Curtis Ordination: an effective strategy for analysis of multivariate ecological data. In Advances in Ecological Research (eds A MacFadyen, ED Ford), pp. 1–55. Academic Press. (doi:10.1016/S0065-2504(08)60168-3)

41. Reas C, Fry B. 2006 Processing: programming for the media arts. AI Soc. 20, 526–538. (doi:10.1007/s00146-006-0050-9)

42. Furneaux B, Bahram M, Rosling A, Yorou NS, Ryberg M. 2021 Long- and short-read metabarcoding technologies reveal similar spatiotemporal structures in fungal communities. Mol. Ecol. Resour. 21, 1833–1849. (doi:10.1111/1755-0998.13387)

43. Xu J. 2017 Fungal DNA barcoding. 6th Int. Barcode Life Conf. 01, 913–932. (doi:10.1139/gen-2016-0046@gen-iblf.issue01)

44. Stielow JB et al. 2015 One fungus, which genes? Development and assessment of universal primers for potential secondary fungal DNA barcodes. Persoonia - Mol. Phylogeny Evol. Fungi 35, 242–263. (doi:10.3767/003158515×689135)

45. Keller S. 2007 Arthropod-pathogenic Entomophthorales: biology, ecology, identification.

46. Hajek AE, Scott KL, Sanchez-Peña SR, Tkaczuk C, Lovett B, Bushley KE. 2025 Annotated checklist of arthropod-pathogenic species in the Entomophthoromycotina (Fungi, Zoopagomycota) in North America. MycoKeys 114, 329–366. (doi:10.3897/mycokeys.114.139257)

47. Jensen AB, Thomsen L, Eilenberg J. 2006 Value of host range, morphological, and genetic characteristics within the Entomophthora muscae species complex. Mycol. Res. 110, 941–950. (doi:10.1016/j.mycres.2006.06.003)

48. Tretter ED, Johnson EM, Wang Y, Kandel P, White MM. 2013 Examining new phylogenetic markers to uncover the evolutionary history of early-diverging fungi: comparing MCM7, TSR1 and rRNA genes for single- and multi-gene analyses of the Kickxellomycotina. Persoonia Mol. Phylogeny Evol. Fungi 30, 106–125. (doi:10.3767/003158513×666394)

49. Manfrino R, Gutierrez A, Ben Gharsa H, Schuster C, López Lastra C, Leclerque A. 2024 Molecular taxonomic characterization and infra-specific diversity of entomopathogenic Beauveria bassiana fungi from Argentina. Fungal Biol. 128, 1800–1805. (doi:10.1016/j.funbio.2024.04.003)

50. Stajich JE, Lovett B, Lee E, Macias AM, Hajek AE, De Bivort BL, Kasson MT, De Fine Licht HH, Elya C. 2023 Signatures of transposon-mediated genome inflation, host specialization, and photoentrainment in Entomophthora muscae and allied entomophthoralean fungi. (doi:10.7554/eLife.92863.1)

51. Edwards S, Naundrup A, Becher PG, De Fine Licht HH. 2025 Patterns of genotype-specific interactions in an obligate host-specific insect pathogenic fungus. J. Evol. Biol. 38, 225–239. (doi:10.1093/jeb/voae149)

52. Steffan BN, Venkatesh N, Keller NP. 2020 Let’s get physical: bacterial-fungal interactions and their consequences in agriculture and health. J. Fungi 6, 243. (doi:10.3390/jof6040243)

53. Partida-Martinez LP, Hertweck C. 2005 Pathogenic fungus harbours endosymbiotic bacteria for toxin production. Nature 437, 884–888. (doi:10.1038/nature03997)

54. Benoit TG, Wilson GR, Pryor N, Bull DL. 1990 Isolation and pathogenicity of Serratia marcescens from adult house flies infected with Entomophthora muscae. J. Invertebr. Pathol. 55, 142–144. (doi:10.1016/0022-2011(90)90047-A)

55. Hammer TJ, Sanders JG, Fierer N. 2019 Not all animals need a microbiome. FEMS Microbiol. Lett. 366, fnz117. (doi:10.1093/femsle/fnz117)

56. Kaur R, Shropshire JD, Cross KL, Leigh B, Mansueto AJ, Stewart V, Bordenstein SR, Bordenstein SR. 2021 Living in the endosymbiotic world of Wolbachia: A centennial review. Cell Host Microbe 29, 879–893. (doi:10.1016/j.chom.2021.03.006)

57. Pawlowska TE, Gaspar ML, Lastovetsky OA, Mondo SJ, Real-Ramirez I, Shakya E, Bonfante P. 2018 Biology of fungi and their bacterial endosymbionts. Annu. Rev. Phytopathol. 56, 289–309. (doi:10.1146/annurev-phyto-080417-045914)

58. Okrasińska A, Bokus A, Duk K, Gęsiorska A, Sokołowska B, Miłobędzka A, Wrzosek M, Pawłowska J. 2021 New endohyphal relationships between Mucoromycota and Burkholderiaceae representatives. Appl. Environ. Microbiol. 87, e02707–20. (doi:10.1128/AEM.02707-20)

59. Łukasik P, Asch M van, Guo H, Ferrari J, Godfray HCJ. 2013 Unrelated facultative endosymbionts protect aphids against a fungal pathogen. Ecol. Lett. 16, 214–218. (doi:10.1111/ele.12031)

60. Bilgo E, Mancini MV, Gnambani JE, Dokpomiwa HAT, Murdochy S, Lovett B, St Leger R, Sinkins SP, Diabate A. 2024 Wolbachia confers protection against the entomopathogenic fungus Metarhizium pingshaense in African Aedes aegypti. Environ. Microbiol. Rep. 16, e13316. (doi:10.1111/1758-2229.13316)

61. Coyle MC, Elya CN, Bronski MJ, Eisen MB. 2024 Entomophthovirus: an insect-derived iflavirus that infects a behavior-manipulating fungal pathogen of dipterans. G3 GenesGenomesGenetics 14, jkae198. (doi:10.1093/g3journal/jkae198)

62. Iwaszkiewicz-Eggebrecht E et al. 2024 High-throughput biodiversity surveying sheds new light on the brightest of insect taxa., 2024.10.25.620209. (doi:10.1101/2024.10.25.620209)

63. Caporaso JG et al. 2012 Ultra-high-throughput microbial community analysis on the Illumina HiSeq and MiSeq platforms. ISME J. 6, 1621–1624. (doi:10.1038/ismej.2012.8)

64. Scholz M, Albanese D, Tuohy K, Donati C, Segata N, Rota-Stabelli O. 2020 Large scale genome reconstructions illuminate Wolbachia evolution. Nat. Commun. 11, 5235. (doi:10.1038/s41467-020-19016-0)

65. Bleidorn C, Gerth M. 2018 A critical re-evaluation of multilocus sequence typing (MLST) efforts in Wolbachia. FEMS Microbiol. Ecol. 94. (doi:10.1093/femsec/fix163)

66. Łukasik P, Kolasa MR. 2024 With a little help from my friends: the roles of microbial symbionts in insect populations and communities. Philos. Trans. R. Soc. Lond. B. Biol. Sci. 379, 20230122. (doi:10.1098/rstb.2023.0122)

67. Lofgren LA, Uehling JK, Branco S, Bruns TD, Martin F, Kennedy PG. 2019 Genome-based estimates of fungal rDNA copy number variation across phylogenetic scales and ecological lifestyles. Mol. Ecol. 28, 721–730. (doi:10.1111/mec.14995)

